# Catalase import into peroxisomes in Arabidopsis

**DOI:** 10.64898/2026.10.10.758186

**Authors:** Zhoubo Hu, Lug Trémulot, Sara Christina Stolze, Hirofumi Nakagami, Michael Hothorn, Roman Ulm

**Affiliations:** Department of Plant Sciences, Section of Biology, Faculty of Sciences, University of Geneva, CH-1211 Geneva 4, Switzerland; Max Planck Institute for Plant Breeding Research, D-50829 Cologne, Germany; Institute of Genetics and Genomics of Geneva (iGE3), University of Geneva, CH-1211 Geneva 4, Switzerland

**Keywords:** catalase, peroxisomal targeting signal (PTS1), PEX5, peroxisomal protein import, reactive oxygen species (ROS), Arabidopsis, co-import

## Abstract

In eukaryotes, catalases are peroxisomal antioxidant enzymes that maintain cellular reactive oxygen species (ROS) homeostasis. Plant catalases lack canonical peroxisomal targeting signals (PTS), and the mechanism underlying their PEX5-dependent peroxisomal import remains poorly understood. Here, we expressed CAT2 with N- and C-terminal GFP fusions in an Arabidopsis *cat2* mutant background and found that only GFP-CAT2 completely restored wild-type function despite comparable catalase activity for both CAT2-GFP and GFP-CAT2. Only GFP-CAT2 localized to peroxisomes, indicating that altered subcellular localization prevented CAT2-GFP functional complementation. Expression of CAT2-GFP in wild-type Arabidopsis and *Nicotiana benthamiana* revealed in vivo interaction with endogenous catalases leading to co-import into peroxisomes. Localization of YFP fused to various CAT2 C-terminal sections identified the CAT2 C-terminal five amino acids as a functional PTS1-like sequence. These findings define a minimal C-terminal targeting motif for plant catalases that is sufficient for their peroxisomal import.

## Introduction

Reactive oxygen species (ROS), including singlet oxygen (^1^O_2_), superoxide anion (O_2•_^−^), hydrogen peroxide (H_2_O_2_), and hydroxyl radicals (•OH), are byproducts of plant energy metabolism and are induced under stress conditions (Halliwell, 2006; Noctor *et al*., 2018; Mittler *et al*., 2022). ROS serve as important signaling molecules, participating in numerous pathways through the oxidation of proteins (Mittler *et al*., 2022; Roy *et al*., 2025; Hu *et al*., 2026). However, excessive accumulation of ROS can exceed plant antioxidant defenses, leading to oxidative stress with major effects including lipid peroxidation (membrane damage), protein oxidation, DNA damage, and disruption of photosynthesis and respiration (Halliwell, 2006; Mittler, 2017). Thus, multiple antioxidant enzymes are involved in ROS scavenging, including catalases (CAT) that are key H_2_O_2_-scavenging enzymes predominantly localized in peroxisomes (Baker *et al*., 2023). Peroxisomes are single membrane–bound organelles present in virtually all eukaryotic cells and are essential for cellular metabolism and redox homeostasis (Kao *et al*., 2018; Pan *et al*., 2020; Kumar *et al*., 2024). Peroxisomal matrix proteins are encoded in the nucleus, synthesized in the cytosol, and imported into peroxisomes via Peroxisomal Targeting Signal 1 (PTS1) or PTS2 that are recognized by the cytosolic receptor PEROXIN 5 (PEX5) and the adaptor protein PEX7, respectively (Skowyra *et al*., 2024). PEX5 and PEX7, the latter anchoring the cargo to a compatible receptor such as PEX5L in animals or PEX18 in yeast, then facilitate transport of folded and oligomeric cargo proteins through the PEX13 translocon (Skowyra *et al*., 2024; Skowyra & Rapoport, 2025). Some peroxisomal matrix proteins without an own intrinsic PTS are translocated into peroxisomes via “piggy-back” co-import with a PTS-containing protein (Gould *et al*., 1989; Glover *et al*., 1994; Kato *et al*., 1996; Lee *et al*., 1997; Nito *et al*., 2002; Cross *et al*., 2016). PTS1 sites are present in the vast majority of peroxisomal matrix proteins at their ultimate carboxy (C)-terminus and are defined as conserved tripeptide motifs, prototypically Ser-Lys-Leu (–SKL) or its functional variants (such as [S/A/C]-[K/R/H]-[L/M], representing a small, basic, and hydrophobic amino acid, respectively) (Brocard & Hartig, 2006; Montilla-Martinez *et al*., 2015; Baker *et al*., 2023). PTS2 sites are present in few peroxisomal matrix proteins near their amino terminus, and consist of less conserved, nonapeptide sequences with the consensus RLx5HL (Montilla-Martinez *et al*., 2015). Whereas PTS1 is generally not cleaved, PTS2 is often cleaved by a protease after peroxisomal import (Schuhmann *et al*., 2008).

Eukaryotic catalases are tetrameric heme-containing enzymes that lack canonical PTS1 and PTS2 sites, yet they directly interact with PEX5 and exhibit PEX5-dependent import into peroxisomes (Chelikani *et al*., 2004; Oshima *et al*., 2008; Otera & Fujiki, 2012). Mammalian catalases possess a C-terminal –KANL sequence that acts as a non-canonical, weak PTS1 (Purdue & Lazarow, 1996; Trelease *et al*., 1996; Shimozawa *et al*., 1999; Otera & Fujiki, 2012). Similarly, fungal catalases possess the C-terminal –SKI/F sequence as a non-canonical, weak PTS1 (Kragler *et al*., 1993; Williams *et al*., 2012). Interestingly, replacement of the catalase C-terminal motif with a canonical PTS1 tripeptide (–SKL) increased catalase binding to PEX5, reduced catalase activity and caused protein aggregation, thus suggesting the endogenous weak, non-canonical signal is important for proper protein folding and tetramerization (Otera & Fujiki, 2012; Williams *et al*., 2012; Al-Hajaya *et al*., 2022).

In Arabidopsis (*Arabidopsis thaliana*), three members of the CAT family show largely distinct expression patterns across tissues (Frugoli *et al*., 1996). CAT1 is mainly expressed in pollen and seeds, CAT2 is in photosynthetic mesophyll cells where it plays a critical role in photorespiration, and CAT3 is in vascular tissues and senescing organs (Hu *et al*., 2010; Baker *et al*., 2023). Like other eukaryotic catalases, plant catalases localize predominantly to peroxisomes; however, their presence has also been documented in nuclei and cytosol (He *et al*., 2021; Al-Hajaya *et al*., 2022; Lin *et al*., 2025; Hu *et al*., 2026). Plant catalases play crucial roles in detoxifying H_2_O_2_, which arises mainly from photosynthesis-associated photorespiration and oxidation of β-fatty acids during seed germination (Brisson *et al*., 1998; Queval *et al*., 2007; Liu *et al*., 2017; Tuzet *et al*., 2019). In Arabidopsis, loss of CAT2 results in lesion formation in leaves under long-day (LD) growth conditions, hypersensitivity to high light and photorespiratory stress, leave upward curling, impaired root elongation, and enhanced tolerance to the catalase inhibitor hydroxyurea (HU) (Vandenabeele *et al*., 2004; Queval *et al*., 2007; Juul *et al*., 2010; Gao *et al*., 2014; Liu *et al*., 2017). Several studies have investigated plant catalase peroxisomal targeting signals. A C-terminal tripeptide (P-S/T-I/M) is conserved in various plant catalases (Mhamdi *et al*., 2012); however, there are conflicting reports regarding the function of the C-terminus in mediating peroxisomal import. The three C-terminal amino acids (–PSI) of cotton catalase were found to be necessary for peroxisomal import in transient expression systems (Mullen *et al*., 1997). By sharp contrast, the C-terminal 10 amino acids of a pumpkin catalase, including its conserved –PSI motif, were not required for the peroxisomal localization of GFP-CAT1 in stably transformed BY-2 tobacco cells and stable transgenic Arabidopsis (Kamigaki *et al*., 2003). Instead, the 13^th^ to 11^th^ ultimate C-terminal terminal amino acids (QKL) have been proposed to function as an internal PTS1 (Kamigaki *et al*., 2003; Fujikawa *et al*., 2019). Recently, expression of an untagged CAT2 isoform lacking the C-terminal 18 amino acids in a *cat2* background functionally complemented the tested phenotypes (Al-Hajaya *et al*., 2022). Aside from catalase C-terminal import signals, several proteins have been reported as required for catalase peroxisomal localization, but the underlying molecular mechanisms remain largely unknown. LON PROTEASE 2 (LON2) is involved in facilitating the import of catalases into peroxisomes (Lingard & Bartel, 2009). LESION SIMULATING DISEASE 1 (LSD1) was proposed to be key for redox-regulated CAT2 compartmentalization between peroxisome, cytosol and nucleus (Lin *et al*., 2025). Moreover, CATALASE-INTERACTING RCC1-LIKE 1 (CAIR1) contributes to proper peroxisomal localization under standard conditions and is required for redox-mediated cytosolic accumulation of CAT2 under oxidative stress conditions (Hu *et al*., 2026).

Despite their essential peroxisomal function, the exact mechanism of catalase translocation into peroxisomes has remained a matter of debate (Baker *et al*., 2023). Here we show that, in Arabidopsis, functional homotetramers of CAT2 and heterotetramers of CAT2 with CAT1 and CAT3 are formed in the cytosol, which are subsequently imported into peroxisomes through interaction of the CAT2 PTS1-like C-terminal five amino acids with the PEX5 receptor.

## Results

### A C-terminal GFP impairs catalase peroxisomal import and *in vivo* function

We generated genomic expression constructs encoding CAT2 with either an N- or C-terminal GFP fusion, designated Pro_CAT2_:GFP-CAT2 and Pro_CAT2_:CAT2-GFP, respectively (Fig. 1A), and stably transformed Arabidopsis *cat2-2* null mutants. We selected two independent CAT2-GFP lines (#32 and #54) and two independent GFP-CAT2 lines (#2 and #6) expressing the respective fusion proteins (Fig. 1B). When grown under LD conditions, the two GFP-CAT2 lines fully restored *cat2* growth, whereas the two CAT2-GFP lines still displayed leaf necrosis, reduced rosette size, flat leaves, and lower fresh weight compared to that in wild type, indicating only partial complementation of *cat2* phenotypes (Fig. 1, C and D). To understand why GFP-CAT2 and CAT2-GFP differed in their functional complementation of *cat2*, we measured extractable catalase activity to test whether C-terminal GFP fusion affected catalytic activity. Catalase extracted from both GFP-CAT2 and CAT2-GFP transgenic lines exhibited catalase enzymatic activity comparable to that in wild type, which was in stark contrast to that in *cat2* (Fig. 1E), indicating that C-terminal GFP fusion to CAT2 does not impair its activity. We then assayed glutathione status as a marker for leaf redox state, which confirmed that, in contrast to GFP-CAT2, CAT2-GFP failed to restore *cat2* redox homeostasis under LD growth conditions (Fig. 1F). Moreover, in agreement with *cat2* complementation capacity, expression levels of the oxidative stress marker genes *GRX480*, *UGT74E2* and *APX1*, as well as the salicylic acid– related cell death marker gene *PR2*, were comparable to wild type in GFP-CAT2 complementation lines, whereas their expression levels were elevated in the CAT2-GFP lines, similar to that in *cat2* (Fig. 1, G to J).

**Fig. 1.**
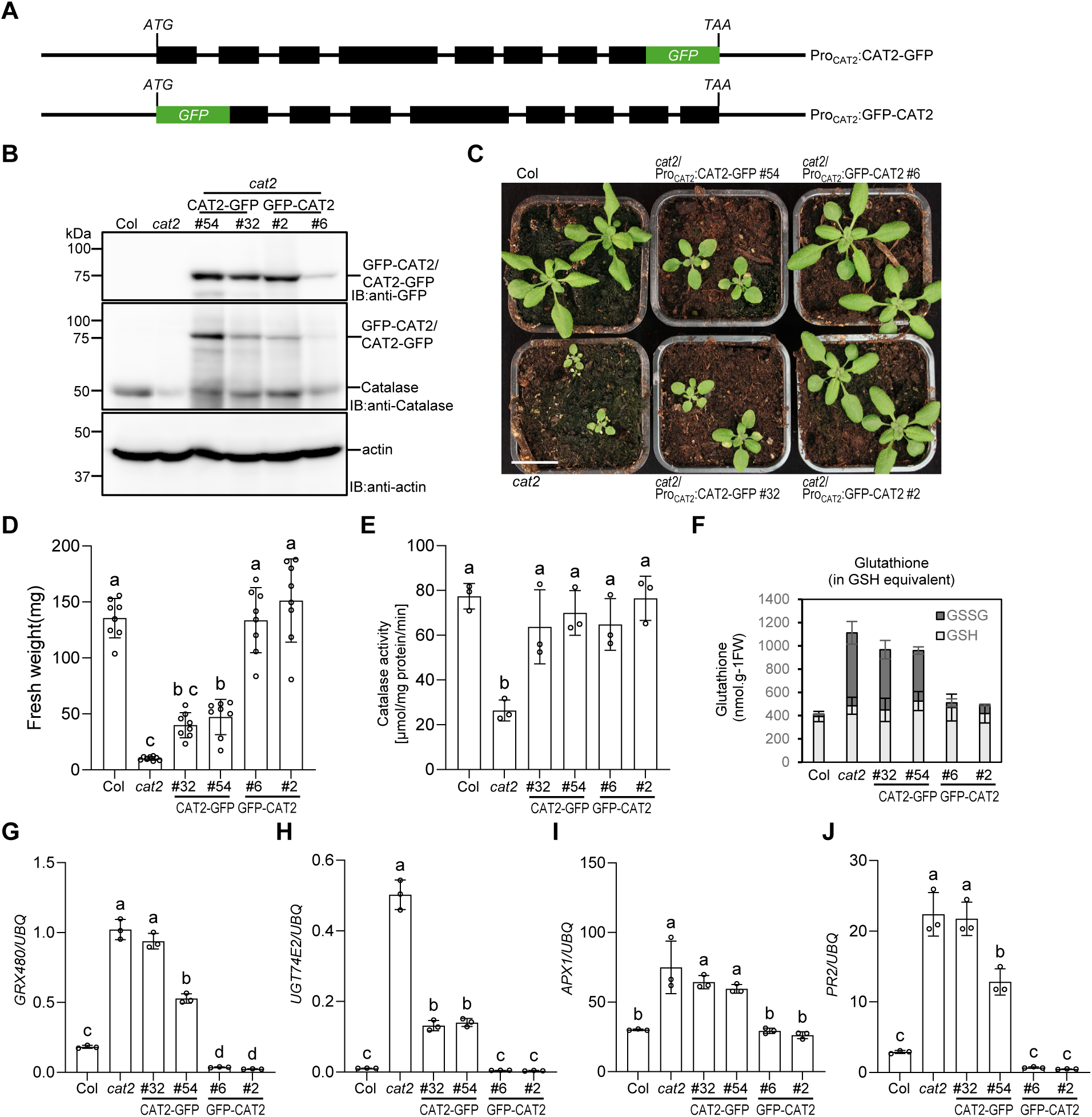
CAT2-GFP only partially rescues the *cat2* phenotype under long-day conditions. (**A**) Schematic representation of the *CAT2* genomic sequence indicating GFP fusion at the N- and C-termini (GFP-CAT2 and CAT2-GFP, respectively). Linker sequences GFP-*YPAFLYKVA*-CAT2 and CAT2-*ASLYKKAGL*-GFP not shown. Black boxes: exons. Black lines: introns, as well as 5′ and 3′ UTRs. (**B**) Immunoblot analysis of catalase protein levels in 10-day-old wild type (Col), *cat2-2* (*cat2*), *cat2-2*/Pro_CAT2_:CAT2-GFP (*cat2* CAT2-GFP, lines #54 and #32), and *cat2-2*/Pro_CAT2_:GFP-CAT2 (*cat2* GFP-CAT2, lines #2 and #6) seedlings, detected using anti-GFP and anti-catalase antibodies. Actin protein levels are shown as loading control. IB, immunoblotting. (**C**) Images of 2-week-old plants grown under long-day (LD) conditions. Scale bar: 2 cm. (**D**) Fresh weight of 2-week-old, LD-grown plants. Independent plant data points and means ± SD (n = 8) are shown. (**E**) Catalase activity as measured in protein extracts from 10-d-old seedlings. Data shown are means ± SD of 3 independent experiments. Shared letters indicate no statistically significant difference between the means (P > 0.05), as determined by two-way ANOVAs followed by Tukey’s test for multiple comparisons. (**F**) Glutathione amount and its redox state were measured in leaves of 3-week-old plants. Dark gray bars represent the reduced form (GSH), and light gray bars represent the oxidized form (GSSG), as GSH equivalent. Data shown are means ± SD of 3 independent experiments. (**G** to **J**) RT-qPCR analysis of *GRX480*, *UGT74E2*, *APX1* and *PR2* gene expression in 2-week-old, LD-grown plants. Independent experiment data points and means ± SD (n = 3) are shown. Shared letters indicate no statistically significant difference between the means (P > 0.05), as determined by two-way ANOVAs followed by Tukey’s test for multiple comparisons.

To further explore the impact of GFP fusion position on CAT2 function, we compared additional stress responses of GFP-CAT2- and CAT2-GFP-complementation lines under specific conditions. Whereas GFP-CAT2 restored HU sensitivity to wild-type levels, CAT2-GFP-expressing lines remained HU tolerant, similar to *cat2* (Fig. 2A). Exogenous supplementation with 0.5% or 1% sucrose, which suppresses ROS production from peroxisomal fatty acid β-oxidation during seedling establishment (Liu *et al*., 2017; Hu *et al*., 2026), significantly reduced HU toxicity (fig. S1), further supporting that HU toxicity is linked to peroxisomal catalase function. We next assessed photosynthetic performance under different oxidative stress conditions by measuring chlorophyll fluorescence to determine *F*v/*F*m, an indicator of the maximum quantum yield of PSII (Baker, 2008; Murchie & Lawson, 2013). Under conditions promoting photorespiration or under high-light stress, GFP-CAT2 plants exhibited *F*v/*F*m values comparable to wild type, whereas CAT2-GFP failed to complement the *cat2* phenotype and exhibited hypersensitivity to oxidative stress (Fig. 2B to D). Together, these results demonstrate that, despite comparable catalase activity, CAT2-GFP provides only partial functional complementation under oxidative stress, whereas GFP-CAT2 fully restores CAT2 function.

**Fig. 2.**
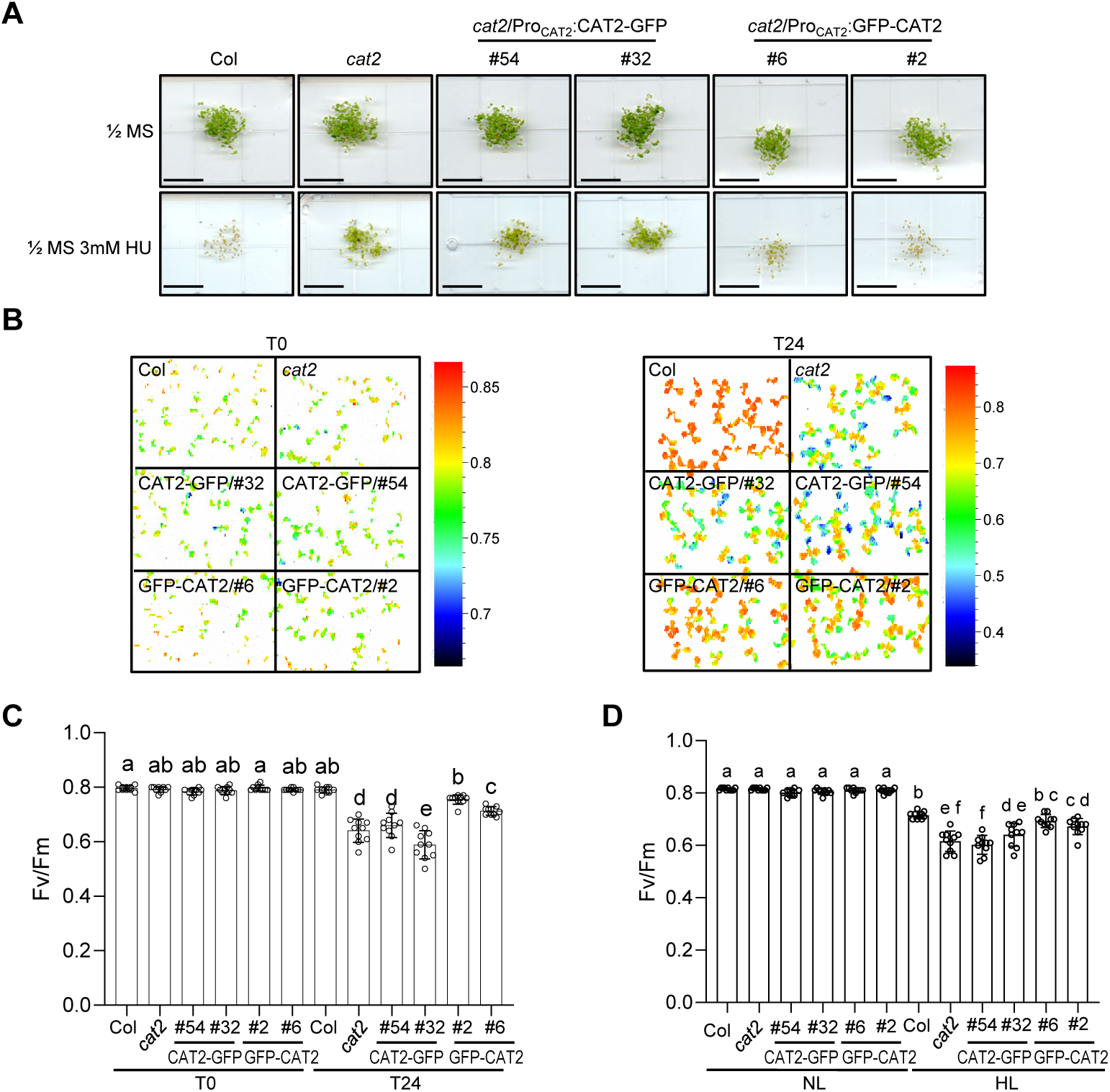
CAT2-GFP does not rescue the *cat2* phenotype under oxidative stress. (**A**) Hydroxyurea (HU) resistance of wild type (Col), *cat2-2* (*cat2*), *cat2-2*/Pro_CAT2_:CAT2-GFP (CAT2-GFP) lines #54 and #32, and *cat2-2*/Pro_CAT2_:GFP-CAT2 (GFP-CAT2) lines #2 and #6. Seedlings were grown on sucrose-free MS medium with or without 3 mM HU. The images were captured 5 days after germination. This experiment was repeated three times with similar results. Scale bar: 1 cm. (**B** and **C**) False-color image representing *F*v/*F*m values (B) and *F*v/*F*m measurements (C) of 10-day-old seedlings before (T0) and after 24 hours (T24) enhanced photorespiration conditions. (**D**) *F*v/*F*m measurements of 10-day-old seedlings before (NL) and after 3-h high light treatment (HL). (C and D) Values of independent measurements and means ± SD are shown (n =10). Shared letters indicate no statistically significant difference between the means (P > 0.05), as determined by two-way ANOVAs followed by Tukey’s test for multiple comparisons.

### C-terminal GFP fusion blocks CAT2-GFP localization to peroxisomes

We hypothesized that C-terminal GFP fusion may impair catalase translocation into peroxisomes. We thus investigated CAT2-GFP subcellular localization in *cat2*/Pro_CAT2_:CAT2-GFP stable transgenic lines. In contrast to the peroxisomal localization of GFP-CAT2 and a GFP-PTS1 peroxisomal marker control, CAT2-GFP was localized to cytosol in leaves, hypocotyls, and roots, with no peroxisomal localization observed (Fig. 3A). We recently showed that enhanced photorespiration and associated elevated ROS prevents CAT2 peroxisomal localization and promotes its cytosolic localization (Hu *et al*., 2026). We thus hypothesized that elevated ROS levels were the cause of CAT2-GFP cytoplasmic localization. This was tested by analyzing CAT2-GFP localization in *cat2*/CAT2-GFP lines under reduced photorespiration conditions achieved by supplementing CO_2_ (fig. S2); however, CAT2-GFP maintained cytosolic localization under high CO_2_ conditions, suggesting that elevated ROS is not preventing CAT2-GFP peroxisomal localization in the *cat2* mutant background.

**Fig. 3.**
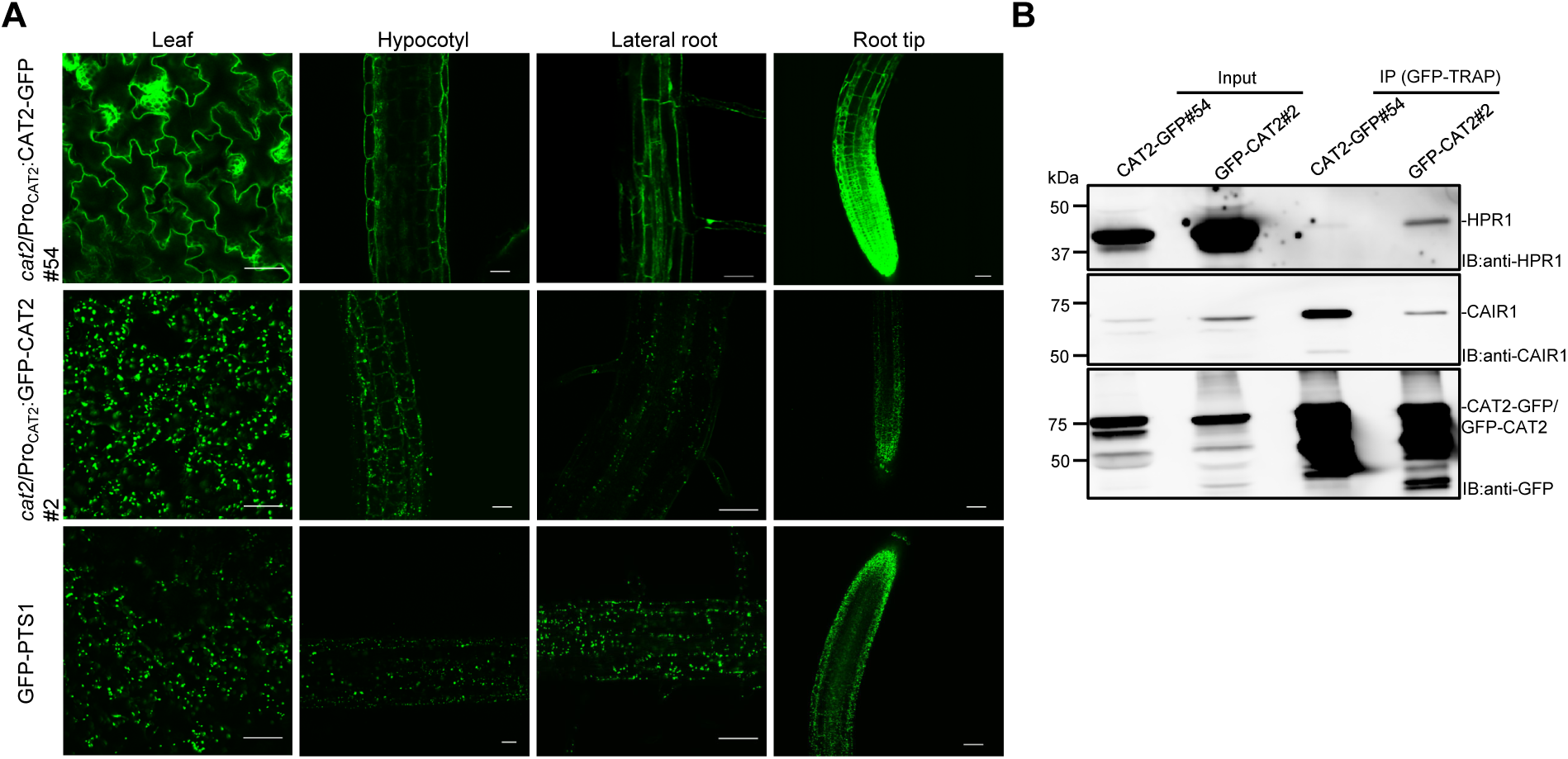
A C-terminal GFP fusion impairs CAT2 localization to peroxisomes. (**A**) GFP signals were detected by confocal microscopy in leaves, hypocotyls, and roots of 10-day-old *cat2-2*/Pro_CAT2_:CAT2-GFP (*cat2*/Pro_CAT2_:CAT2-GFP #54), *cat2-2*/Pro_CAT2_:GFP-CAT2 (*cat2*/Pro_CAT2_:GFP-CAT2 #2) and Pro_35S_:GFP-PTS1 (GFP-PTS1) seedlings. Scale bars: 20 μm. Images in (A) are representative of three experimental repeats. (**B**) Co-IP of HPR1 and CAIR1 with N- and C-terminally GFP-tagged CAT2. Proteins were extracted from 10-day-old *cat2-2*/Pro_CAT2_:CAT2-GFP (CAT2-GFP#54) and *cat2-2*/Pro_CAT2_:GFP-CAT2 (GFP-CAT2#2) seedlings and detected using anti-GFP, anti-HPR1 and anti-CAIR1 antibodies. IB, immunoblotting.

To identify and compare CAT2 interactors from different subcellular compartments, we used 10-day-old *cat2*/Pro_CAT2_:GFP-CAT2 and *cat2*/Pro_CAT2_:CAT2-GFP seedlings for affinity purification coupled to mass spectrometry (AP-MS) using anti-GFP complex purification. As controls, protein extracts from seedlings of wild type, Pro_35S_:StrepII-3xHA-YFP (herein “YFP”) and a peroxisomal marker line Pro_35S_:GFP-PTS1 (GFP-PTS1) were processed in parallel to reveal non-specific binding proteins. Several previously identified interactors (CAT1, CAT3, NCA1, CAIR1 and HPR1) were detected from both GFP-CAT2 and CAT2-GFP purification (table S1) (Hackenberg *et al*., 2013; Li *et al*., 2015; Liu *et al*., 2019; Hu *et al*., 2026). Notably, PEX5 co-purified with GFP-CAT2 but not with CAT2-GFP (Table 1 and table S1). We compared our AP-MS performed with seedling extracts with our previous GFP-CAT2 purification from three-week-old plants (Hu *et al*., 2026) (table S2). Many CAT2 interactors were uniquely identified in either mature plant or seedling samples (table S2, fig. S3, A, C and D). We further focused on 28 proteins identified to co-purify with GFP-CAT2 independent of plant age or growth conditions (fig. S3, A and B, and table S2). We then compared AP-MS of *cat2*/CAT2-GFP and *cat2*/GFP-CAT2 with the reasoning that their specific subcellular localization would highlight cytoplasmic and peroxisomal CAT2 interactors, respectively. Indeed, for *cat2*/CAT2-GFP, there were reduced interactions with peroxisomal proteins (incl. HPR1, KAT5, and ACX4) and enhanced interactions with cytosolic proteins (incl. CAIR1, HSP60) (Table 1). In agreement, co-IP of the peroxisomal localized HPR1 with GFP-CAT2 was enhanced in comparison to that with CAT2-GFP, whereas the inverse was observed for the cytosolic localized CAIR1 (Fig. 3B). Altogether, we conclude that a C-terminal GFP fusion prevents CAT2 translocation into peroxisomes, likely due to impaired PEX5 interaction and peroxisomal import.

**Table 1:**
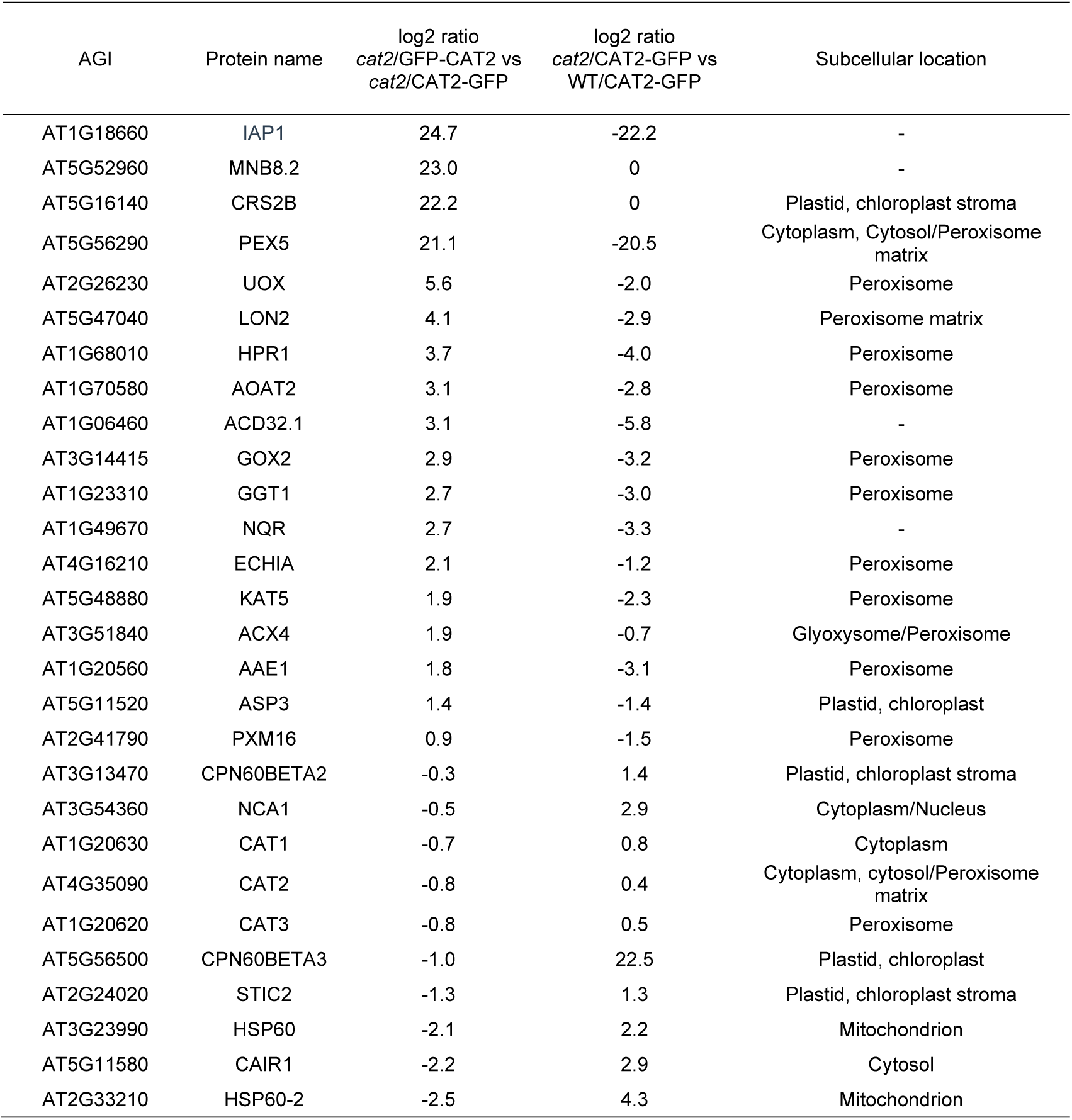
Comparison of AP-MS–identified interactors of GFP-CAT2 and CAT2-GFP. *cat2-2*/Pro_CAT2_:GFP-CAT2 line #2 (*cat2*/GFP-CAT2), *cat2-2*/Pro_CAT2_:CAT2-GFP line #54 (*cat2*/CAT2-GFP) and Col-0/Pro_CAT2_:CAT2-GFP line #4 (Col/CAT2-GFP) lines. Subcellular location is specified according to UniProt Annotation, except for AT5G11580/CAIR1 which is according to Hu et al., 2026. Corresponding data in table S1. AGI, Arabidopsis Genome Initiative locus identifier.

### CAT2 forms functional oligomers prior to import into peroxisomes

Further investigation into CAT2-GFP localization dynamics in a wild-type background indicated that oligomerization with endogenous CAT2 allows co-import of CAT2-GFP into peroxisomes. In sharp contrast to that seen in the *cat2* null mutant background, CAT2-GFP showed peroxisomal localization in Col/Pro_CAT2_:CAT2-GFP (Fig. 4A). Furthermore, unlike CAT2-GFP cytosolic localization in the parental *cat2*/Pro_CAT2_:CAT2-GFP line, CAT2-GFP localized to peroxisomes in the F1 generation of a backcross to wild type, but not in that for the control cross to *cat2* (fig. S4A). Moreover, in the F1 of a genetic cross between *cat2*/Pro_CAT2_:CAT2-GFP and *cat2*/Pro_CAT2_:GFP-CAT2, GFP signals were only found in peroxisomes (fig. S4B), thus further indicating that CAT2-GFP interaction with GFP-CAT2 allows CAT2-GFP co-translocation into peroxisomes through a “piggyback” co-import mechanism. Of note, both CAT2-GFP and GFP-CAT2 localized to peroxisomes during transient expression assays in *N. benthamiana*, and exhibited overlapping signals with peroxisomal marker mCherry-PTS1 (fig. S4B). This suggests that proper peroxisomal localization of CAT2-GFP is also possible via oligomerization with *N. benthamiana* catalase.

**Fig. 4.**
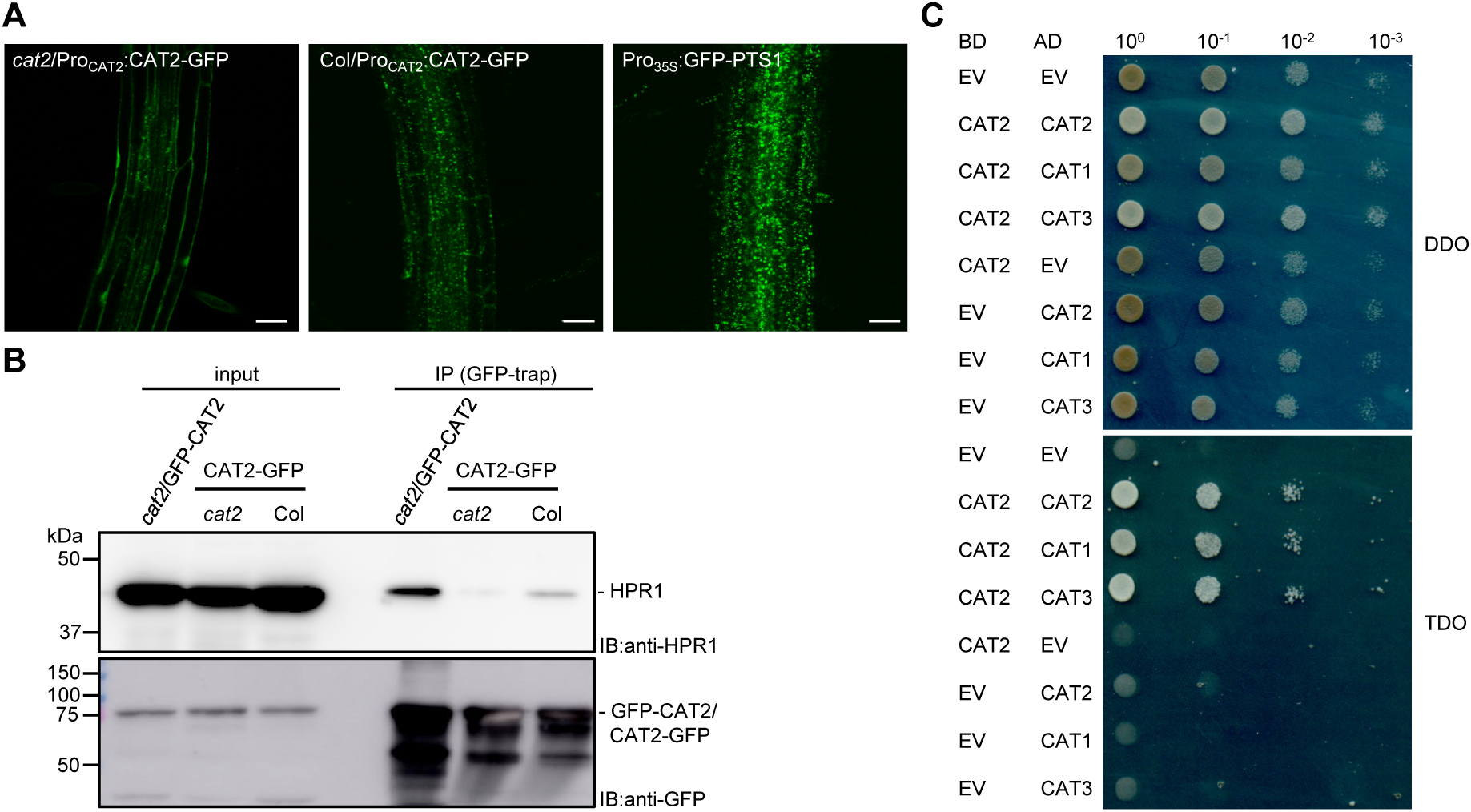
CAT2-GFP localizes to peroxisomes in the presence of endogenous catalase. (**A**) CAT2-GFP localizes to peroxisomes in the stable transgenic line Col/Pro_CAT2_:CAT2-GFP but not in the *cat2* mutant background. GFP fluorescence was examined by confocal microscopy in the roots of 10-day-old *cat2-2*/Pro_CAT2_:CAT2-GFP, Col/Pro_CAT2_:CAT2-GFP, and Pro_35S_:GFP-PTS1 seedlings. Scale bar: 20 μm. Images in (A) are representative of three replicate experiments. (**B**) Co-IP of peroxisomal HPR1 with CAT2-GFP is enhanced in wild-type Col background compared to *cat2* background. Proteins were extracted from 10-day-old *cat2*/Pro_CAT2_:GFP-CAT2 (*cat2*/GFP-CAT2), *cat2*/Pro_CAT2_:CAT2-GFP (*cat2*/CAT2-GFP), and Col/Pro_CAT2_:CAT2-GFP (Col/CAT2-GFP) seedlings and detected using anti-HPR1 and anti-GFP antibodies. IB, immunoblotting. (**C**) Interaction of CAT2 with CAT1, CAT2, and CAT3 in a yeast two-hybrid growth assay. Tenfold serial dilutions of transformed yeast spotted on SD/−Trp/−Leu (DDO; nonselective for interaction) and SD/−Trp/−Leu/−His (TDO; selective) plates. AD: Activation domain. BD: DNA binding domain, EV: empty vector.

Similarly to purifications of CAT2-GFP and GFP-CAT2 expressed in the *cat2* background, AP-MS of CAT2-GFP from Col/Pro_CAT2_:CAT2-GFP identified CAT1, CAT3, NCA1 and CAIR1 (table S1). Interestingly, however, PEX5, which was not purified with CAT2-GFP in the *cat2* background, was co-purified with CAT2-GFP expressed in the wild-type background (table S1). We also found that known peroxisomal interactors of catalases were enriched, whereas cytosol interactors were reduced in CAT2-GFP AP-MS performed using Col/ Pro_CAT2_:CAT2-GFP compared to using *cat2*/ Pro_CAT2_:CAT2-GFP (Table 1), which reflected the differential CAT2-GFP localization in the two backgrounds. In agreement, co-IP of HPR1 with CAT2-GFP was enhanced when the latter was expressed in wild type rather than in *cat2* (Fig. 4B). CAT1 and CAT3 co-purified with GFP-CAT2 and CAT2-GFP (table S1) (Hu *et al*., 2026), and CAT1, CAT2 and CAT3 interacted with each other in yeast two-hybrid assays (Fig. 4C). Altogether we conclude that CAT2-GFP localizes to peroxisomes when expressed in a wild type background due to interactions with endogenous catalases, particularly CAT2, thus allowing CAT2-GFP to co-opt a functional PTS via cytosolically assembled homo- and heterotetrameric catalase complexes.

### C-terminal VRPSI^488–492^ pentapeptide from CAT2 functions as PTS1 sufficient for peroxisomal localization

The C-terminal amino acid sequences of Arabidopsis CAT1, CAT2, and CAT3 are generally well conserved, although the terminal tripeptide PSI in CAT2 and CAT3 diverged to PNF in CAT1 (Fig. 5A). We thus focused on the major Arabidopsis catalase in leaves, CAT2. To define the functional PTS of CAT2 and to eliminate interference from interactions with endogenous catalases, we fused CAT2 C-terminal peptides to the C-terminus of YFP and assessed their subcellular localization using transient expression assays in *N. benthamiana* leaves (Fig. 5B). YFP alone localized to the cytosol, whereas fusion of the canonical PTS1 tripeptide SKL was sufficient to target YFP-SKL to peroxisomes (Fig. 5C). Similarly, the ultimate C-terminal peptide of both fungal and human catalase, respectively, directed YFP-SKF and YFP-KANL fusion proteins into peroxisomes, demonstrating that, despite not being canonical PTS1 peptides, the PTS of fungal and human catalases are functional in plants (Fig. 5C). We then tested the CAT2 C-terminal tripeptide PSI (CAT2^490–492^) fused to YFP, which localized to the cytosol unlike the peroxisomal marker mCherry-PTS1, indicating that PSI alone is insufficient for peroxisomal targeting (Fig. 5C). We next examined whether a larger C-terminal region of CAT2, particularly the region including the internal tripeptide QKL that has been suggested to function as internal PTS1 (Kamigaki *et al*., 2003; Fujikawa *et al*., 2019), could confer peroxisomal localization. Fusion of the last 13 amino acids, CAT2^480–492^, indeed successfully directed YFP to peroxisomes (Fig. 5C). However, the QKL tripeptide alone failed to target YFP to peroxisomes and YFP-CAT2^483–492^ missing the QKL tripeptide maintained peroxisomal localization, indicating that the C-terminal 10 amino acids of CAT2 are sufficient (Fig. 5C). Although the C-terminal PSI was not sufficient, it was required for peroxisomal targeting, as the C-terminally truncated YFP-CAT2^480–489^ as well as a PSI-AAA mutated version, YFP-CAT2^483–492;PSI-AAA^, failed to localize to peroxisomes (Fig. 5C). Finally, systematically truncating the ultimate C-terminal 10 amino acids of CAT2 revealed that the last 5 amino acids, –VRPSI, were sufficient for peroxisomal localization of YFP-CAT2^488–492^, whereas YFP-CAT2^489–492^ remained cytosolic (Fig. 5D). We conclude that the C-terminal pentapeptide VRPSI constitutes a functional, non-canonical PTS1 in Arabidopsis CAT2, which is sufficient for import into peroxisomes.

**Fig. 5.**
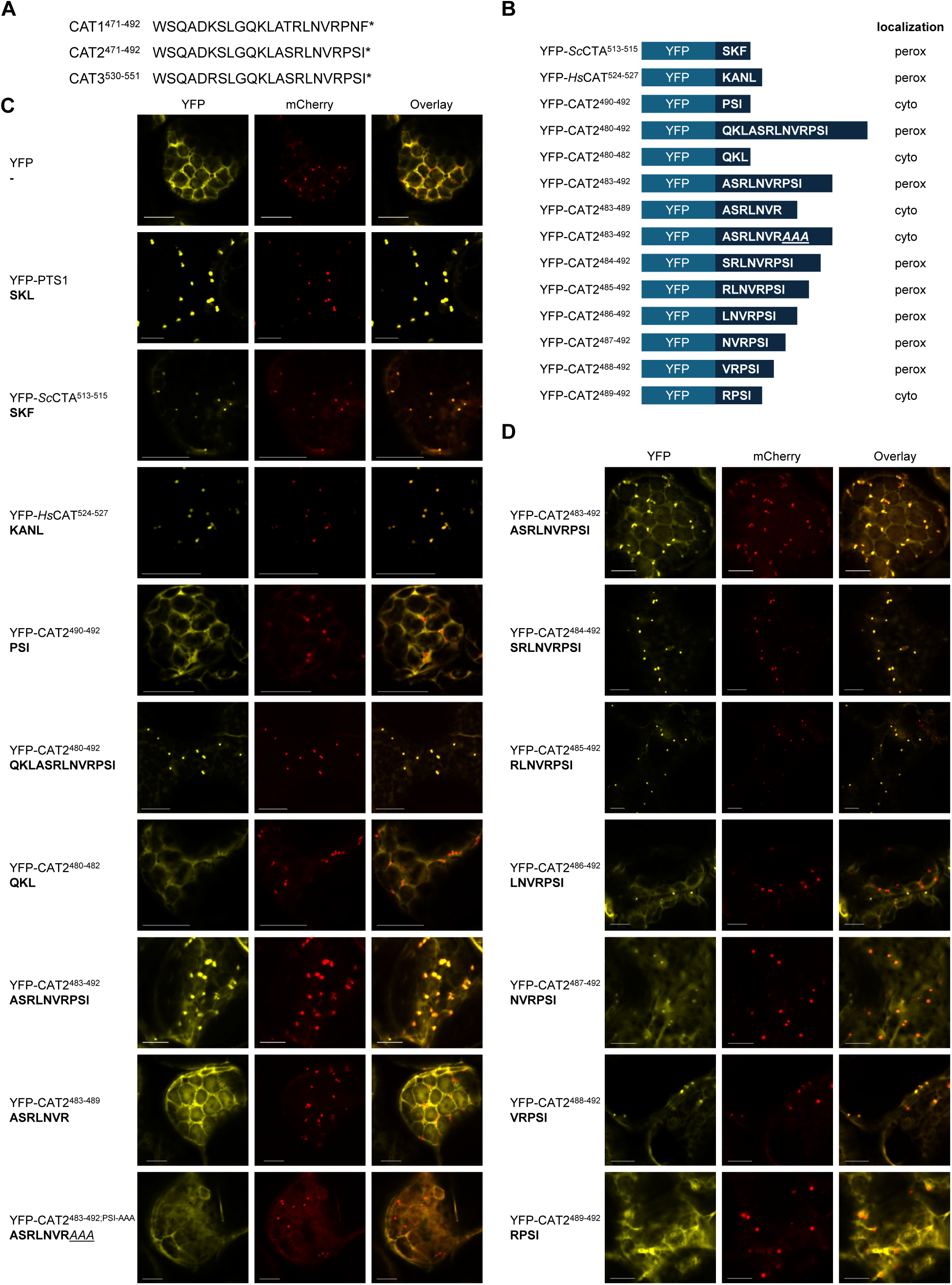
The C-terminal 5 amino acids of Arabidopsis CAT2 are both necessary and sufficient for peroxisomal localization. (**A**) Alignment of the C-terminal 20 amino acids from Arabidopsis CAT1, CAT2, and CAT3. Asterisks (*) indicate translation terminates after the preceding amino acid. (**B**) Schematic representation of various CAT2 C-terminal amino acids fused C-terminally to YFP and their intracellular localization when transiently expressed in *N. benthamiana* leaves. ScCTA: *Saccharomyces cerevisiae* CTA (NP_010542.1); HsCAT: *Homo sapiens* CAT (NP_001743). Perox, peroxisome; cyto, cytosol. (**C** and **D**) Representative confocal microscopy images of YFP localization upon transient expression of YFP with the indicated C-terminally fused peptide sequences in *N. benthamiana* leaves. Images of co-transformed Pro_35S_:mCherry-PTS1 (SKL tripeptide) are shown as a positive control for peroxisomal localization. Scale bars: 10 μm.

To structurally characterize the interaction of CAT2 with PEX5, we generated AlphaFold3 complex models of the full-length enzyme with the tetratricopeptide repeats (TPR) domain of the receptor (Fig. 6A). In the resulting model, the unstructured C-terminus of CAT2 binds to the canonical peptide binding groove of the PEX5 TRP domain (Fig. 6B). Strikingly, despite the sequence differences, CAT2^487–492^ binds PEX5 in a similar fashion as previously described for classical PTS1 sequences (Gatto *et al*., 2000; Stanley *et al*., 2006; Sampathkumar *et al*., 2008) (Fig. 6C), suggesting that sequence variations are well tolerated by the receptor. The ultimate C-terminal residue, however, needs to be hydrophobic and is a major determinant of the interaction, consistent with the mutational analysis. Indeed, the C-terminal isoleucine in CAT2 adopts a similar conformation as the leucine in the SKL motif of a canonical PTS1 (Fig. 6C). We speculate that the CAT2 C-terminus binds to PEX5 more weakly compared to that of a canonical PTS1 motif, and thus a longer peptide that includes Val-488 mediates a more robust interaction for peroxisomal import. Indeed, Val-488 binds to a hydrophobic groove in PEX5 in our AlphaFold model (Fig. 6B). Notably, Arabidopsis CAT1 and CAT3, as well as human catalase, may share a highly similar binding mode (Fig. 6D). Overall, we conclude that plant catalases use their C-terminal –VRPSI pentapeptides as PTS1 for PEX5 interaction and peroxisomal import.

**Fig. 6.**
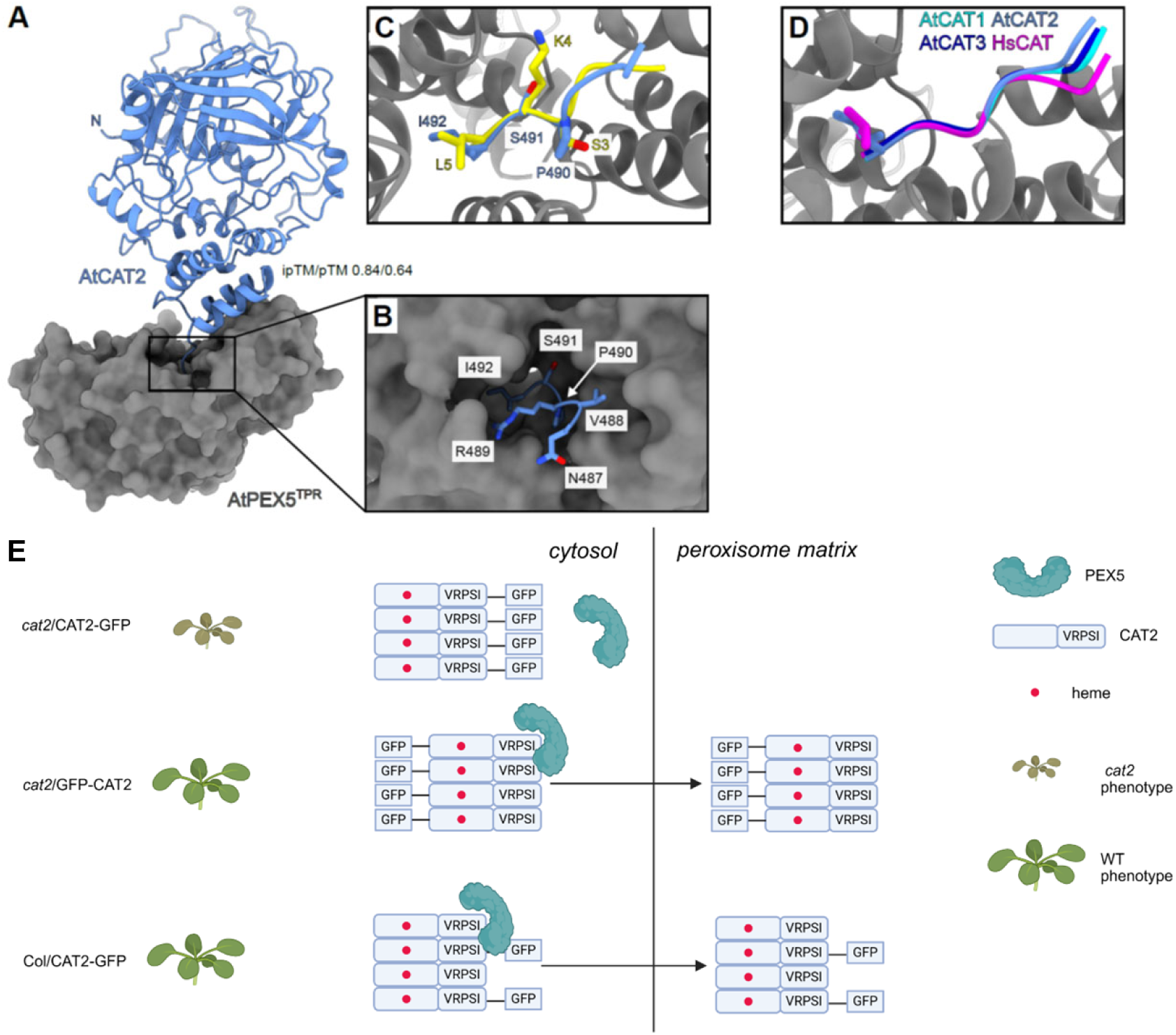
Arabidopsis catalase import into peroxisomes using a C-terminal PTS1-like sequence. The C-termini of plant and human catalases adopt a PTS1-like binding mode. (**A**) Alphafold3 model (predicted template modeling scores ipTM = 0.84, pTM = 0.64) of the Arabidopsis CAT2–PEX5 complex. The C-terminus of CAT2 (depicted as a blue ribbon) binds to the canonical PTS1 binding groove in the PEX5 TPR domain (residues 442-728, molecular surface, in gray). (**B**) Close-up view of the peptide binding groove in PEX5 bound to the C-terminus of CAT2. CAT2 interacting residues 487–492 are highlighted in the bonding representation. (**C**) Structural superposition of the CAT2–PEX5 complex model with the previously reported crystal structure of the human HsPEX5–PTS1 complex (PDB-ID 1FCH (*52*); root mean square deviation r.m.s.d. is ∼0.8 Å comparing 260 corresponding C_α_ atoms in the TPR domain, shown in gray). Arabidopsis CAT2 residues 482–492 are depicted as a ribbon (in blue) and the PTS1 (SKL) peptide is shown in yellow. The C-terminal three amino acids in both peptides are highlighted in bonds representation. (**D**) Structural superposition of the Alphafold3 predicted complexes of Arabidopsis CAT1–PEX5 (in cyan, ipTM=0.72, pTM=0.62), CAT2–PEX5 (in blue), and CAT3–PEX5 (in dark blue, ipTM=0.82, pTM=0.63), together with human HsCAT–HsPEX5 (in magenta, ipTM=0.78, pTM=0.69). The complex models align with an r.m.s.d of ∼0.7 Å comparing 421 C_α_ atoms. (**E**) Model illustrating peroxisomal import capacity of GFP-CAT2 and CAT2-GFP. GFP-CAT2 forms a functional, heme-containing catalase tetramer and retains its native C-terminal five amino acids (–VRPSI) as PTS1, enabling interaction with PEX5 and subsequent import into peroxisomes. As a result, GFP-CAT2 complements the *cat2* mutant phenotype. By contrast, CAT2-GFP also assembles into a functional, heme-containing tetramers in the cytosol, but its C-terminal PTS1 is masked by the C-terminal GFP fusion, preventing interaction with PEX5. Consequently, CAT2-GFP cannot be imported into peroxisomes and thus does not fully complement the *cat2* mutant phenotype. In the wild-type (Col) background, however, CAT2-GFP can assemble into heterotetramers with endogenous catalase subunits. These mixed complexes retain an exposed PTS1 provided by the endogenous catalase, enabling PEX5-mediated import of the entire complex into peroxisomes. This model was made with Biorender (https://www.biorender.com/).

## Discussion

Catalase, the primary H_2_O_2_-decomposing enzyme in peroxisomes, is essential for maintaining cellular ROS homeostasis. Despite its fundamental role, the mechanism by which catalase is imported into peroxisomes is disputed to date (Mullen *et al*., 1997; Kamigaki *et al*., 2003; Fujikawa *et al*., 2019; Al-Hajaya *et al*., 2022; Baker *et al*., 2023). Here, we investigated the peroxisomal import mechanism of Arabidopsis CAT2 by comparing the activity, localization, function, and interactome of CAT2-GFP and GFP-CAT2. Our results indicate that CAT2 assembles into functional homo- and/or heterotetramers in the cytosol prior to peroxisomal import. Furthermore, we demonstrate that the C-terminal five amino acids of CAT2 are sufficient for peroxisomal targeting, identifying them as a functional PTS1 that mediates PEX5-dependent import.

Catalases are predominantly localized to peroxisomes in eukaryotes (Mhamdi *et al*., 2012), but the timing and location of their maturation have remained controversial. Pulse-chase studies in mammalian cells suggested that catalases are imported as monomers and subsequently acquire heme and assemble into tetramers within the peroxisome (Lazarow & de Duve, 1973; Freitas *et al*., 2011). By contrast, fibroblasts from patients with Zellweger syndrome, which lack functional peroxisomes, contain enzymatically active tetrameric catalase in the cytosol (Brul *et al*., 1988). Together with the observation that recombinant catalase expressed in *E. coli* is catalytically active (Li *et al*., 2015; Turek *et al*., 2020), these findings imply that neither peroxisomal localization nor import is strictly required for catalase folding, heme incorporation, or enzymatic activity. Our results support this latter model. Although CAT2-GFP localized predominantly to the cytosol in the *cat2* mutant background, extractable catalase activity remained comparable to that of wild type, indicating that CAT2 can form an active enzyme outside the peroxisome.

Several lines of evidence support the idea that catalase oligomerization precedes peroxisomal import. Despite the CAT2-GFP C-terminal GFP fusion masking the endogenous targeting signal, CAT2-GFP localized efficiently to peroxisomes when expressed in a wild-type background or transiently in *N. benthamiana*. This result is most readily explained by a piggyback co-import mechanism with endogenous catalases that maintain an exposed C-terminal targeting sequence, thus indicating that a functional PTS1 is not essential for every catalase subunit within the tetramer. Consistent with this model, our yeast two-hybrid and AP-MS analyses indicated that CAT2 interacts with both itself and with CAT1 and CAT3, suggesting the formation of both homo- and heterotetramers, despite their partially distinct expression patterns (Frugoli *et al*., 1996). Similar mechanisms have been reported in mammalian cells, where a C-terminally truncated catalase is imported into peroxisomes only in the presence of wild-type catalase (Otera & Fujiki, 2012). Piggyback co-import has also been demonstrated for several other oligomeric peroxisomal proteins, including thiolase, isocitrate lyase, and chloramphenicol acetyltransferase, whose PTS1-deficient forms are co-imported with PTS1-containing partners (Glover *et al*., 1994; Lee *et al*., 1997).

Nevertheless, CAT2-GFP only partially complemented the physiological defects of the *cat2* mutant, which may be explained by limited peroxisomal import through piggyback co-import with endogenous CAT1 or CAT3. These observations indicate that catalase maturation—including folding, heme incorporation, and oligomerization—occurs in the cytosol (Fig. 6E), whereas proper peroxisomal localization is required for its full physiological function. Whether catalytically active cytosolic catalase contributes directly to ROS homeostasis remains unclear. Interestingly, mammalian catalase accumulates in the cytosol under oxidative stress (Fujiki & Bassik, 2021), and recent studies similarly reported stress-induced cytosolic accumulation of *Arabidopsis* CAT2 during enhanced photorespiration and elevated ROS production (Lin *et al*., 2025; Hu *et al*., 2026). Together with our findings, this raises the possibility that cytosolic CAT2 has a physiological function beyond its established role in peroxisomes.

Our data further clarify the long-standing ambiguity surrounding peroxisomal targeting signals of plant catalases. Human catalase contains the C-terminal tetrapeptide –KANL that is both necessary and sufficient for peroxisomal targeting (Purdue & Lazarow, 1996). Interestingly, C-terminal fusion of –KANL also directed YFP to peroxisomes in *N. benthamiana*. The ultimate C-terminal sequences of plant catalases are highly divergent from those of mammal and fungal catalases; however, their ultimate ∼20 amino acids are remarkably conserved across plant species (Mhamdi *et al*., 2012). Previous studies investigating the importance of this region reached conflicting conclusions (Al-Hajaya *et al*., 2022; Baker *et al*., 2023), likely because endogenous catalases enabled piggyback co-import in both stable and transient expression systems. By separating direct targeting from co-import, we demonstrate that the ultimate C-terminal pentapeptide of CAT2 (–VRPSI) is sufficient to target a YFP reporter to peroxisomes in *N. benthamiana*. These findings establish –VRPSI as a functional PTS1-like sequence and suggest that plant catalases evolved a sequence-divergent, yet fully functional, peroxisomal targeting signal.

PEX5 is the cytosolic receptor for PTS1-containing proteins and typically recognizes canonical PTS1 motifs through its C-terminal TPR domain (Gatto *et al*., 2000; Stanley *et al*., 2006; Sampathkumar *et al*., 2008). Accordingly, mutations in PEX5 disrupt catalase localization in mammals and plants (Piekarska *et al*., 2006; Lingard *et al*., 2009; Otera & Fujiki, 2012; Dubreuil *et al*., 2020). However, previous studies proposed that catalases from mammals, yeast, and plants interact preferentially with the N-terminal region of PEX5 rather than the canonical TPR domain (Oshima *et al*., 2008; Freitas *et al*., 2011; Rymer *et al*., 2018). By contrast, our AlphaFold3 structural models predict that plant, yeast and human catalases all engage PEX5 in a binding mode closely resembling that of canonical PTS1-containing cargo proteins. Despite substantial divergence in their ultimate C-terminal sequences, all catalases are predicted to occupy the same PEX5 binding pocket. These models suggest that the hydrophobic character of the terminal residues, rather than strict sequence conservation, is the principal determinant of PEX5 recognition. This provides a structural explanation for how catalases with highly diverse C-terminal sequences can nevertheless be imported through a conserved PEX5-dependent mechanism.

In summary, our findings support a model in which plant catalase folds, incorporates heme, and assembles into functional homo- and heterotetramers in the cytosol before being imported into peroxisomes. Moreover, we identify the CAT2 C-terminal pentapeptide –VRPSI as a functional PTS1-like sequence and provide evidence that it is recognized by PEX5 through a canonical PTS1-binding mechanism despite its non-canonical sequence.

## Materials and methods

### Plant materials and growth conditions

*cat2-2* (SALK_057998) (Queval *et al*., 2007), *cat2-2/*Pro_CAT2_:GFP-CAT2 (Hu *et al*., 2026), and Pro_35S_:GFP-PTS1 (Zolman *et al*., 2000) are in the Columbia accession (Col-0).

For experiments under aseptic conditions, Arabidopsis seeds were surface sterilized with chlorine gas and sown on half-strength Murashige & Skoog basal salt (MS) medium containing 0.5% w/v phytagel (Sigma) without sucrose. After 2 d of stratification at 4°C, seedlings were grown under LD conditions (16-h/8-h light/dark) in a standard growth chamber (MLR-350, Sanyo) with 66 µmol m^-2^ s^-1^ white light (measured with a LI-250 light meter; LI-COR Biosciences).

For rosette phenotypes, seeds were sown on soil and stratified for 2 d at 4°C before transfer to LD conditions with 120 µmol m^-2^ s^-1^ white light in a GroBank growth chamber (CLF Plant Climatics).

For photorespiratory stress, Petri dishes, containing plants grown on half-strength MS medium without sucrose for 7 d, were sealed with three layers of Parafilm M (Amcor) to restrict the ambient air influx and then exposed to continuous light (200 µmol m^-2^ s^-1^ white light) to prevent the confounding effects of night-time respiration on gas homeostasis (Waszczak *et al*., 2016).

For high-light treatment, Petri dishes, containing plants grown on half-strength MS medium with 1% w/v sucrose for 7 d under LD conditions in a standard growth chamber (MLR-350, Sanyo) with 66 µmol m^-2^ s^-1^ white light, were transferred to high-light conditions (1,000 µmol m^-2^ s^-1^) using a JBeamBio LED panel for a 3-h treatment.

### Construct preparation

To construct Pro_CAT2_:CAT2-GFP, three fragments were first amplified and cloned into entry vectors using MultiSite Gateway Recombination Cloning Technology (primers listed in table S1): (1) a fragment containing the 5ʹ regulatory sequence (943 bp) of *CAT2* and the *CAT2* genomic sequence (2276 bp) that was amplified and cloned into pDONR P4-P1R, (2) the coding sequence of GFP including stop codon that was amplified from N-SpR-GFP plasmid (Hu *et al*., 2019) and cloned into pDONR221, and (3) a fragment containing *CAT2* 3ʹ regulatory sequence (924 bp) that was amplified and cloned into pDONR P2R-P3. The entry vectors pDONR P4-P1R, pDONR221 and pDONR P2R-P3 carrying the above respective fragments were cloned into pFR7m34GW (Kalmbach *et al*., 2023) by MultiSite Gateway cloning technology to generate Pro_CAT2_:CAT2-GFP genomic clones.

To construct Pro_35S_:YFP-PSI and Pro_35S_:YFP-13AA, the fragments YFP-PSI and YFP-13AA were amplified from pGWB541 (Nakagawa *et al*., 2007) (primers listed in table S1) and cloned into entry vector pDONR207 using the Gateway system (Invitrogen). Then pDONR207-YFP-PSI and pDONR207-YFP-13AA were cloned into pGWB502 (Nakagawa *et al*., 2007). To construct Pro_35S_:mCherry-PTS1(SKL), the fragment mCherry-PTS1(SKL) was amplified from N-KmR-mCherry (Hu *et al*., 2019) (primers listed in table S1) and cloned into entry vector pDONR207 using the Gateway system (Invitrogen). Then pDONR207-mCherry-PTS1 was cloned into pGWB502 (Nakagawa *et al*., 2007).

To construct AD-CAT1, AD-CAT2 and AD-CAT3, pDONR207-CAT1, pDONR207-CAT2 and pDONR207-CAT3 were cloned into pGADT7-GW (Lu *et al*., 2010).

### Catalase activity measurements

Catalase activity was measured as previously described, according to the original protocol (Aebi, 1984). Briefly, approximately 50 mg leaf material was ground in liquid nitrogen and total protein was extracted with 1.5 mL 0.1 M PBS buffer (pH 7.5). The homogenates were centrifuged for 10 min at 14000 rpm at 4°C, and the supernatants were used for the assay. The protein concentration was measured with Bradford assay.

### Glutathione amount and redox state quantification

Leaves of 3-week-old plants grown in soil were used for the quantification of glutathione and its redox state, according to a published protocol (Noctor & Mhamdi, 2022). Briefly, extraction was performed in liquid nitrogen and 0.2 M HCl using a mortar and pestle. Neutralization was then performed with 0.2 M NaOH. Glutathione concentration was determined by measuring the rate of increase in absorbance at 412 nm (A_412_) during reduction of 5,5’-dithio-bis-(2-nitrobenzoic acid) (DTNB) in the presence of NADPH and glutathione reductase. For each sample and standard curve point, three technical replicates are performed in a 96-well plate (Greiner Bio-One) using a TECAN Spark plate reader.

### Immunoblotting analysis and co-immunoprecipitation

For immunoblotting, total proteins were extracted in lysis buffer (50 mM Tris-HCl [pH 7.5], 10% v/v glycerol, 1 mM EDTA, 150 mM NaCl, 1 mM PMSF, 5mM DTT, and 1 tablet of cOmplete^™^, EDTA-free Protease Inhibitor Cocktail [Sigma-Aldrich]). Total protein was quantified with Quick Start™ Bradford Protein Assay (Bio-Rad) according to the manufacturer’s instructions. 25 μg of protein per sample was separated by SDS-PAGE and transferred onto PVDF membranes with iBlot3 (Invitrogen). GFP-tagged proteins, CAIR1, HPR1, and actin were detected using Living Colors^®^ GFP Monoclonal Antibody (1:5,000; Clontech, 632381), anti-CAIR1(1:1,000) (Hu *et al*., 2026), anti-HPR1 (1:10,000; Agrisera, AS111797) and anti-actin antibodies (1:10,000; Sigma, A0480), respectively, in TBST buffer containing 5% w/v milk, as described previously (Hu *et al*., 2026). Catalases were detected using anti-catalase (1:1,000; Agrisera, AS09501) in PBST buffer containing 5% w/v milk. Detection was performed using horseradish peroxidase (HRP)-conjugated goat anti-rabbit (1:10,000; Vector PI-1000) and anti-mouse (1:10,000; ThermoFisher) secondary antibodies, followed by ECL Select Western Blotting Detection Reagent (GE Healthcare) and imaging in an Amersham Imager 680 camera system (GE Healthcare).

For co-immunoprecipitation, 1 g of plant tissue was ground to a fine powder in liquid nitrogen and homogenized in 2 mL of lysis buffer (50 mM Tris-HCl [pH 7.5], 10% v/v glycerol, 1 mM EDTA, 150 mM NaCl, 1 mM PMSF, 5mM DTT, 0.1% v/v Triton X-100, and cOmplete^™^, EDTA-free Protease Inhibitor Cocktail [Sigma-Aldrich]). After centrifugation at 12,000 rpm (17,000 × g) for 20 min at 4°C, the supernatant was collected and protein concentration determined by the Bradford assay. The same amount of total protein was incubated with 30 μL of ChromoTek GFP-Trap® Agarose (Proteintech) for 1 h at 4°C with gentle shaking. The beads were pelleted at 500 rpm and washed three times with 2 mL of washing buffer (50 mM Tris-HCl [pH 7.5], 300 mM NaCl). Bound proteins were eluted by boiling the beads for 5 min in 40 μL of 1× Laemmli buffer (63 mM Tris-HCl [pH 6.8], 10% v/v glycerol, 2% v/v SDS, 0.1% v/v 2-mercaptoethanol, 0.0005% w/v bromophenol blue). Subsequently, 20 μL of the supernatant was analyzed by western blotting as described above.

Immunoblotting experiments and co-IP assays were repeated three times with similar results.

### Affinity purification–mass spectrometry (AP-MS)

*cat2-2*/Pro_CAT2_:CAT2-GFP line #54, *cat2-2*/Pro_CAT2_:GFP-CAT2 line #2, and Col-0/Pro_CAT2_:CAT2-GFP line #4 were grown for 10 d on sucrose-free half-strength MS medium in a standard growth chamber (MLR-350, Sanyo) under white light (66 µmol m⁻² s⁻¹). Approximately 6 g of seedlings were harvested, ground into a fine powder in liquid nitrogen, and homogenized in 12 mL of lysis buffer (50 mM Tris-HCl [pH 7.5], 10% v/v glycerol, 1 mM EDTA, 150 mM NaCl, 1 mM PMSF, 5 mM DTT, 0.1% v/v Triton X-100, and 1 tablet of cOmplete^™^, EDTA-free Protease Inhibitor Cocktail [Sigma-Aldrich]). The extracts were centrifuged at 7,200 *g* for 20 min at 4°C. For each sample, triplicate technical subsamples were taken of the supernatant, each comprising 8 mg of total protein, which were adjusted to an equal volume of 5 mL in lysis buffer and used for immunoprecipitation. The extracts were incubated for 1 h at 4°C with 40 µL GFP-Trap^®^ agarose resin (ChromoTek) that had been prewashed three times with 1.5 mL lysis buffer and resuspended in 100 μL lysis buffer before addition to the protein samples. The resins were pelleted at 500 rpm at 4°C and washed four times with 5 mL washing buffer (50 mM Tris–HCl [pH 7.5], 300 mM NaCl). Then, the resin was resuspended in 1 mL washing buffer, transferred to 1.5-mL Eppendorf tubes, and pelleted again. As controls, triplicate protein extracts were prepared in parallel from wild type (Col), Pro_35S_:StrepII-3xHA-YFP (Lapin *et al*., 2019) and Pro_35S_:GFP-PTS1 (Zolman *et al*., 2000).

Sample preparation and LC-MS/MS data acquisition: enriched proteins were submitted to an on-bead digestion using trypsin. In brief, dry beads were re-dissolved in 25 µL digestion buffer 1 (50 mM Tris, pH 7.5, 2 M urea, 1 mM DTT, 5 ng/µL trypsin) and incubated for 30 min at 30°C in a Thermomixer with 400 rpm. Next, beads were pelleted and the supernatant was transferred to a fresh tube. Digestion buffer 2 (50 mM Tris, pH 7.5, 2M urea, 5 mM CAA) was added to the beads and mixed, after which the beads were pelleted, the supernatant collected and combined with the previous collected supernatant. The combined supernatants were then incubated o/n at 32°C in a Thermomixer set to 400 rpm, with samples protected from light during incubation. The digestion was stopped by adding 1 µL TFA and desalted with C18 Empore disk membranes according to the StageTip protocol (Rappsilber *et al*., 2003).

Dried peptides were re-dissolved in 2% v/v ACN, 0.1% v/v TFA (10 µL) for analysis. Samples were analyzed using an Ultimate 3000 RSLC nano (Thermo Fisher) coupled to an Orbitrap Exploris 480 mass spectrometer equipped with a field asymmetric ion mobility separation (FAIMS) Pro interface (Thermo Fisher). Peptides were pre-concentrated on an Acclaim PepMap 100 pre-column (75 µM x 2 cm, C18, 3 µM, 100 Å, Thermo Fisher) using the loading pump and buffer A** (water, 0.1% v/v TFA) with a flow of 7 µl/min for 5 min. Peptides were separated on 16-cm frit-less silica emitters (New Objective, 75 µm inner diameter), packed in-house with reversed-phase ReproSil-Pur C18 AQ 1.9-µm resin (Dr. Maisch). Peptides were loaded on the column and eluted for 130 min using a segmented linear gradient of 5% to 95% solvent B (0 min : 5%B; 0–5 min -> 5%B; 5–65 min -> 20%B; 65–90 min ->35%B; 90–100 min -> 55%; 100–105 min ->95%, 105–115 min ->95%, 115–115.1 min -> 5%, 115.1–130 min ->5%) (solvent A 0% ACN, 0.1% FA; solvent B 80% ACN, 0.1% FA) at a flow rate of 300 nL/min. Mass spectra were acquired in data-dependent acquisition mode with a TOP_S method using a cycle time of 2 seconds. For FAIMS, two compensation voltages (-45 and -60) were applied, the cycle times for both compensation voltages were set to 1 s. MS spectra were acquired in the Orbitrap analyzer with a mass range of 320–1200 m/z at a resolution of 60,000 FWHM and a normalized AGC target of 300%. Precursors were filtered using the MIPS option (MIPS mode = peptide), the intensity threshold was set to 5000, precursors were selected with an isolation window of 1.6 m/z. HCD fragmentation was performed at a normalized collision energy of 30%. MS/MS spectra were acquired with a target value of 75% ions at a resolution of 15,000 FWHM, at an injection time of 120 ms and a fixed first mass of m/z 120. Peptides with a charge of +1, greater than 6, or with unassigned charge state were excluded from fragmentation for MS2.

Data analysis: raw data were processed using MaxQuant software (version 1.6.3.4, http://www.maxquant.org/) (Cox & Mann, 2008) with label-free quantification (LFQ) and iBAQ enabled (Tyanova *et al*., 2016). MS/MS spectra were searched by the Andromeda search engine against a combined database containing the sequences from *A. thaliana* (TAIR10_pep_20101214; ftp://ftp.arabidopsis.org/home/tair/Proteins/TAIR10_protein_lists/), and sequences of 248 common contaminant proteins and decoy sequences. Trypsin specificity was required and a maximum of two missed cleavages allowed. Minimal peptide length was set to seven amino acids. Carbamidomethylation of cysteine residues was set as fixed, oxidation of methionine and protein N-terminal acetylation as variable modifications. The match between runs option was enabled. Peptide-spectrum-matches and proteins were retained if they were below a false discovery rate (FDR) of 1% in both cases. Statistical analysis of the MaxLFQ values was carried out using Perseus (version 1.5.8.5, http://www.maxquant.org/). Quantified proteins were filtered for reverse hits and hits “identified by site” and MaxLFQ values were log2 transformed. After grouping samples by condition only those proteins were retained for the subsequent analysis that had two valid values in one of the conditions. Two sample T-tests were performed using a permutation-based FDR of 5%. Alternatively, quantified proteins were grouped by condition and only those hits were retained that had three valid values in one of the conditions. Missing values were imputed from a normal distribution (1.8 downshift, separately for each column). The Perseus output was exported and further processed using Excel. Candidates with LFQ intensity value of at least 2 replicates > 18, log2 ratio > 1 and peptides number >2 compared to wild type Col, StrepII-3xHA-YFP and GFP-PTS1 AP-MS negative controls were retained (table S2B). The GFP-CAT2 AP-MS raw data were retrieved from (Hu *et al*., 2026) and analyzed in parallel (table S2A).

The MS data have been deposited in the ProteomeXchange Consortium via the PRIDE partner repository with the dataset identifier PXD080797.

### Yeast two-hybrid assays

Yeast two-hybrid assays were performed after transforming the Y2Hgold strain (Takara) using the lithium-acetate protocol (Gietz, 2014). For growth assays, -His selective medium for interactions (SD/−Trp/−Leu/–His; Formedium) was used.

### Confocal image

The localization of GFP- and mCherry-tagged proteins in Arabidopsis seedlings or *N. benthamiana* was captured by a Leica TCS SP8 confocal microscope (Leica, Bensheim, Germany). GFP was excited with Argon laser at 488 nm, and the emitted fluorescence was detected between 493 and 550 nm. mCherry was excited at 561 nm and the emitted fluorescence was detected between 576 and 632 nm. Line intensity profiles of selected regions of interest (ROI) were generated using the Image J software (Rueden *et al*., 2017) and the data were exported to Excel.

### Transient expression in *Nicotiana benthamiana*

To transiently express proteins in *Nicotiana benthamiana*, constructs were introduced into Agrobacterium strain GV3101 and cultures were co-infiltrated into young leaves of *N. benthamiana* together with P19 gene silencing suppressor protein (Lombardi *et al*., 2009).

### RNA extraction and reverse transcription quantitative PCR (RT-qPCR)

Total RNA from plant tissues was isolated using the Plant RNeasy kit (Qiagen), including DNase treatment, following the manufacturer’s instructions. cDNA synthesis was carried out using the Taqman Reverse Transcription Reagent kit (Thermo Fisher Scientific). Each RT-qPCR reaction contained cDNA equivalent to 3 ng of RNA, synthesized with a 1:1 mixture of oligo(dT) primers and random hexamers. RT-qPCR was performed using PowerUp SYBR Green Master Mix (Thermo Fisher Scientific) on a QuantStudio™ 5 Real-Time PCR System (Thermo Fisher Scientific). Gene expression levels were calculated using the ΔΔCT method, with *UBQ5* (*UBIQUITIN5*, AT3G62250) and *ACT2* (*ACTIN2*, AT3G18780) as reference genes (Czechowski *et al*., 2005). Each experiment included three independent biological replicates.

### *F*v/*F*m measurement

The maximum quantum efficiency of PSII (*F*v/*F*m) was assayed by measuring plant chlorophyll fluorescence using a PSI (Photon Systems Instruments) FluorCam 800 MF, as previously described (Leonardelli *et al*., 2024). Before measurements, plates were incubated in darkness for 7 min, which was confirmed to be sufficient to reach steady state under the plant growth conditions used (Murchie & Lawson, 2013). Maximum fluorescence in the dark (*F*m) was measured using saturating pulses of 2000 µmol m^-2^ s^-1^ (white light, 400–720 nm) for 960 ms, followed by orange-red light (620 nm) detection pulses (10 µs). *F*v/*F*m was calculated as *F*v/*F*m = (*F*m − *F*o)/*F*m, where *F*m is the maximal fluorescence and *F*o is the minimal fluorescence in the dark-adapted state.

### CAT–PEX5 complex predictions

Complex models of Arabidopsis catalases–PEX5 and human HsCAT–HsPEX5 complexes were generated using Alphafold3 (Abramson *et al*., 2024) and displayed and analyzed using ChimeraX (Meng *et al*., 2023).

## Acknowledgments

We would like to thank Emilie Demarsy for helpful comments and critically reading the manuscript, Christophe Belin (University of Perpignan, France) for kindly providing seeds of the peroxisome maker line GFP-PTS1, Lothar Kalmbach (University of Neuchâtel, Switzerland) for plasmid pLOK180_pFR7m34GW, and Anne Harzen (Max Planck Institute for Plant Breeding Research) for MS sample preparation. This work was supported by the Max Planck Society, the University of Geneva and, in part, the Swiss National Science Foundation (grant 310030_207716 to R.U.).

## Competing interests

The authors declare that they have no competing interests associated with this work.

## Author contributions

ZH performed all the experiments, except the measurement of reduced glutathione which was performed by LT. S.C.S and H.N. performed and analyzed LC-MS/MS data. M.H. contributed the structural analysis. ZH and RU designed the research, analyzed the data, and wrote the article, with input from all authors.

## Data availability

Sequence data from this work can be found in the Arabidopsis Genome Initiative or GenBank/EMBL databases under the following accession numbers: CAT1 (AT1G20630), CAT2 (AT4G35090), CAT3 (AT1G20620), HPR1 (AT1G68010), CAIR1 (AT5G11580), PEX5 (AT5G56290), LON2 (AT5G47040), PR1 (AT2G14610), PR2 (AT3G57260), GRX480 (AT1G28480), APX1 (AT1G07890) and UGT74E2 (AT1G05680). UniProt IDs for HsCAT and HsPEX5 are P04040 and P50542, respectively.

---

**Supplemental Figure S1.**
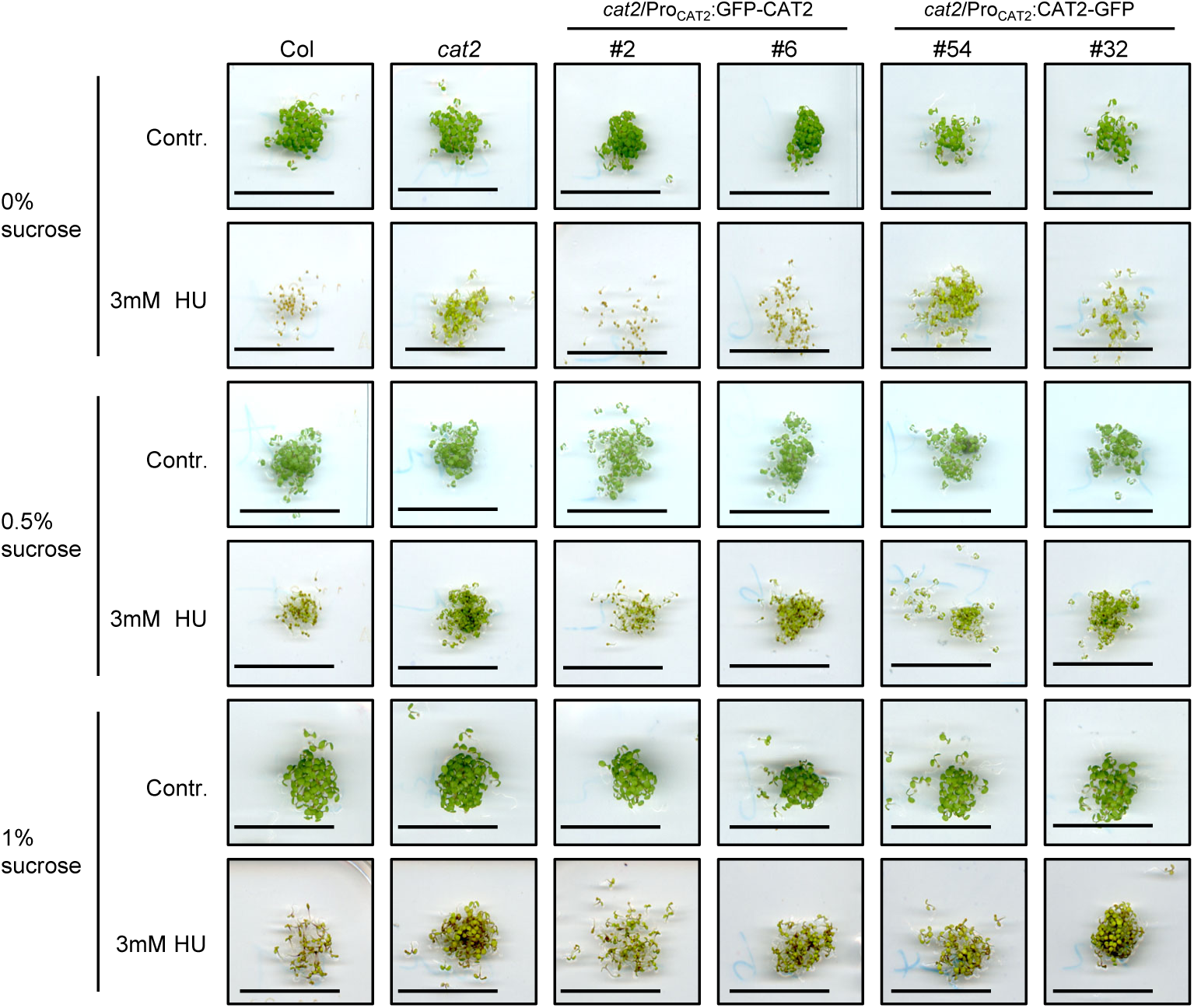
GFP-CAT2 but not CAT2-GFP restores HU sensitivity when expressed in *cat2*, whereas exogenous sucrose suppresses wild-type HU sensitivity. Seeds of wild-type (Col), *cat2-2* (*cat2*), *cat2-2*/Pro_CAT2_:GFP-CAT2 (lines #2 and #6), and *cat2-2*/Pro_CAT2_:CAT2-GFP (lines #54 and #32), were plated on ½ MS medium without (Contr.; control) or with 3 mM HU, in presence of 0.5%, 1% or no (0%) sucrose, as indicated. The images were captured 5 days after germination. Scale bar: 2 cm. This experiment was repeated three times with similar results.

**Supplemental Figure S2.**
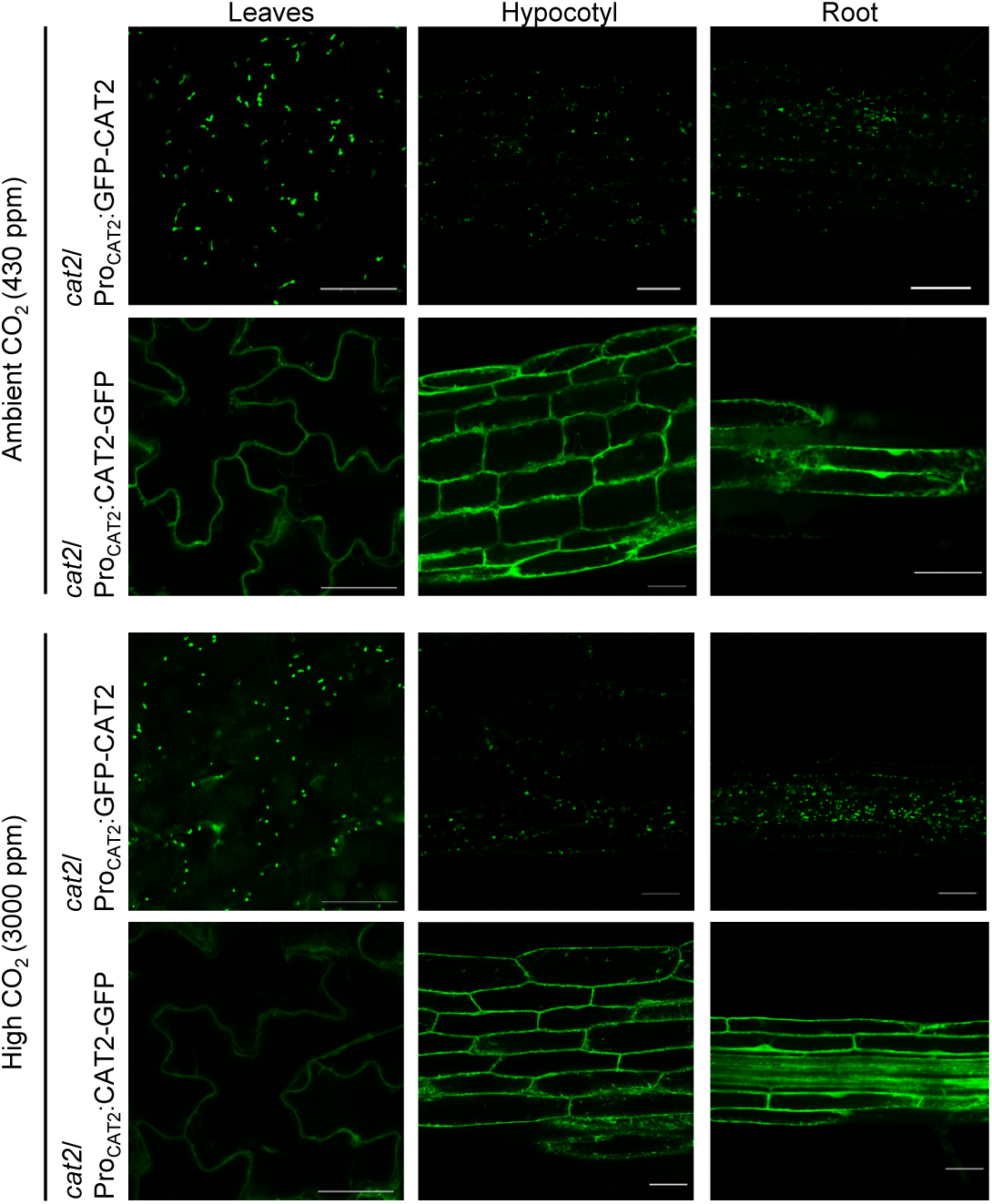
Subcellular localization of GFP-CAT2 and CAT-GFP under elevated CO_2_. Reduced oxidative stress due to reduced photorespiration through CO_2_ supplementation did not alter the cytosolic subcellular localization of CAT2-GFP. GFP signals were observed in 8-day-old *cat2-2*/Pro_CAT2_:GFP-CAT2 (line #6) and *cat2-2*/Pro_CAT2_:CAT2-GFP (line #54) seedlings grown under long-day conditions with high CO_2_ supplementation (3000 ppm) or under ambient CO_2_ (430 ppm; regular air). Scale bar: 10 μm. Representative images from three independent experiments are shown.

**Supplemental Figure S3.**
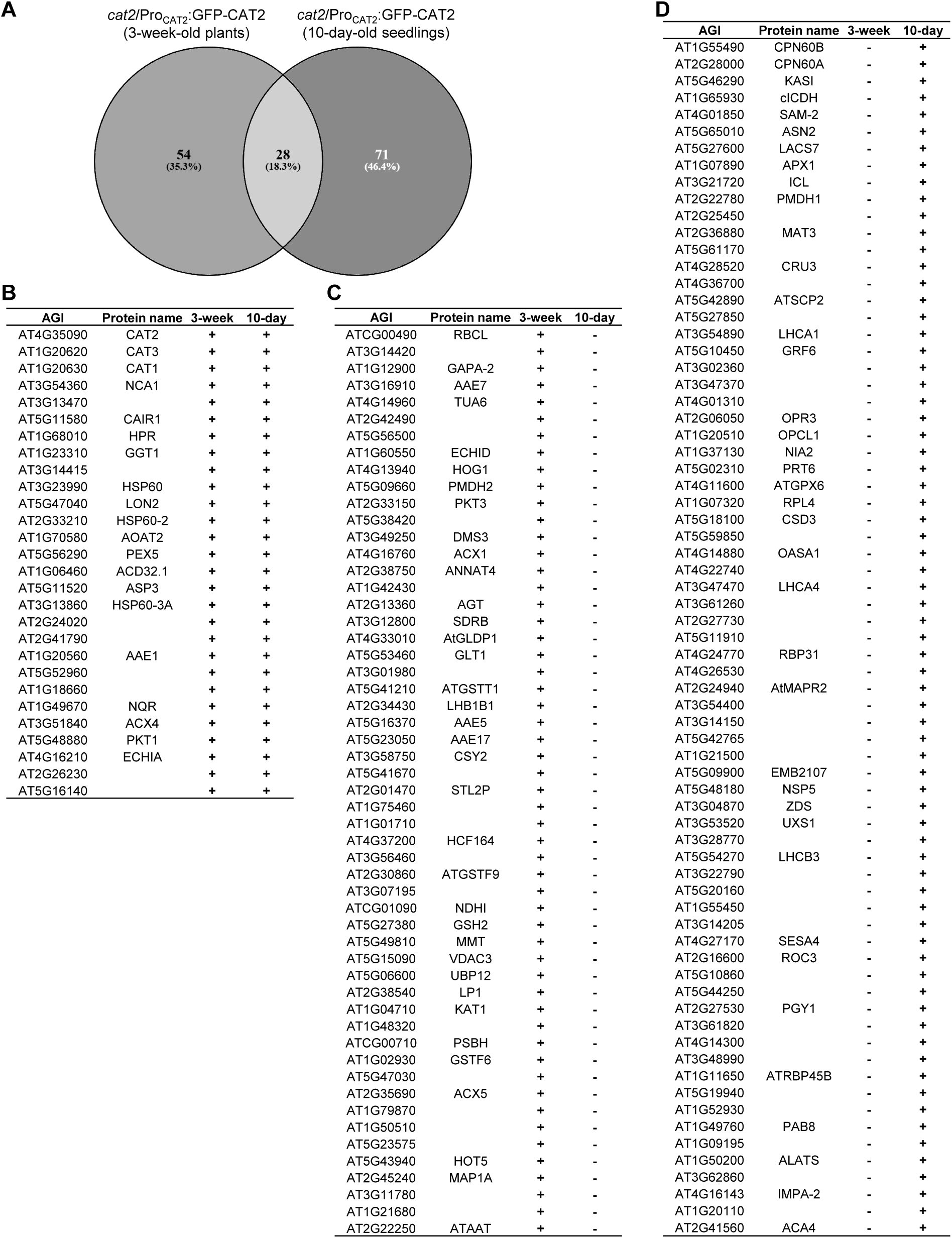
Comparison of proteins identified by GFP-CAT2 AP-MS from 3-week-old plants versus 10-day-old seedlings. **(A)** Venn diagram showing the overlap of candidate GFP-CAT2 interactors identified in 3-week-old plants and 10-day-old seedlings of *cat2-2*/Pro_CAT2_:GFP-CAT2. Data available in table S2. **(B to -D)** Corresponding protein lists; +, identified in AP-MS, -, not identified.

**Supplemental Figure S4.**
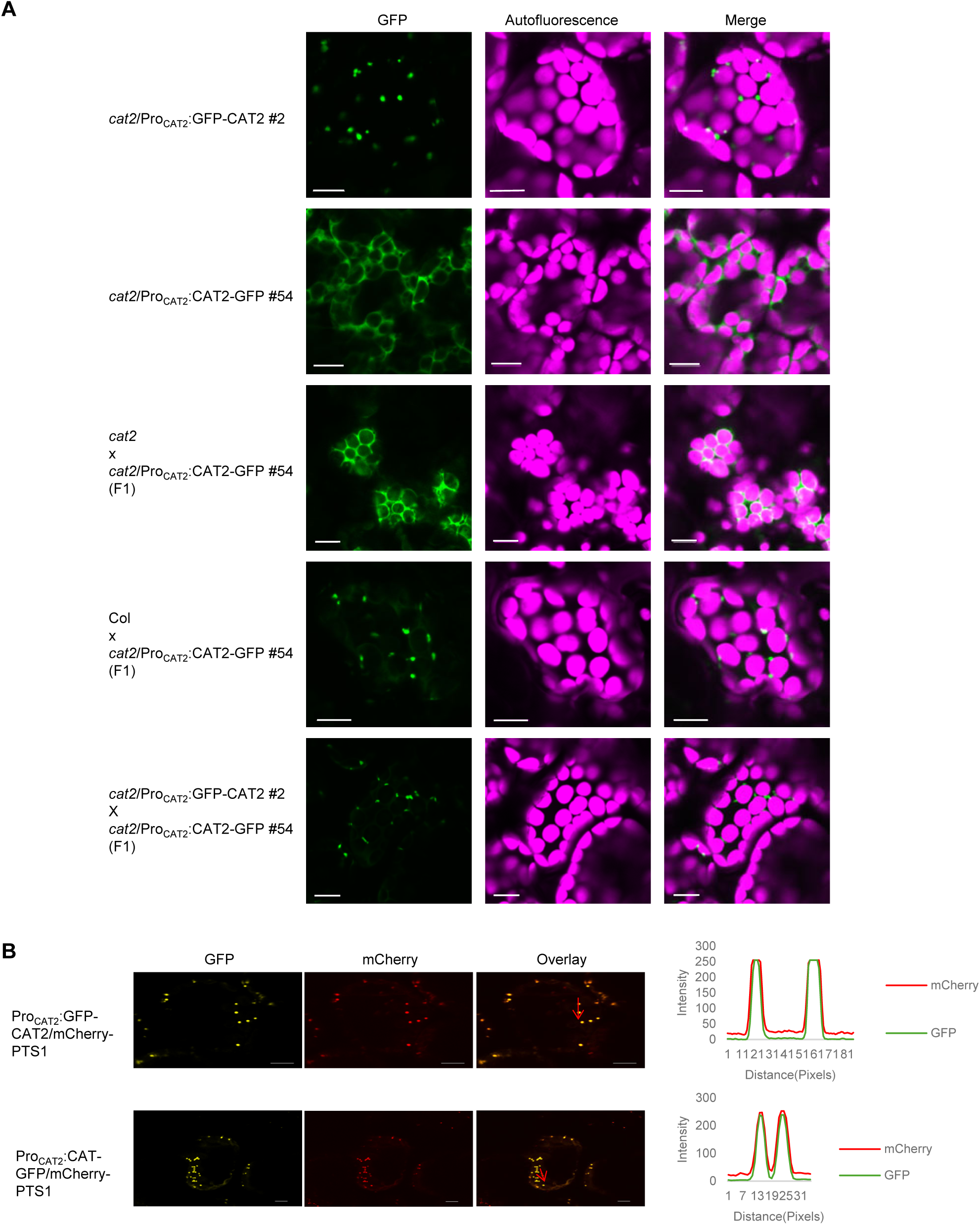
Endogenous CAT2 co-imports CAT2-GFP into peroxisomes. **(A)** The *cat2-2*/Pro_CAT2_:CAT2-GFP line #54 was crossed with wild type Col, *cat2-2*, and *cat2-2*/Pro_CAT2_:GFP-CAT2 line #2, then GFP signals were examined by confocal microscopy in 8-day-old seedlings of the F1 generation, in comparison to the two parental transgenic lines. Scale bars: 10 µm. **(B)** Both GFP-CAT2 and CAT2-GFP showed peroxisomal localization in transiently transformed *N. benthamiana* leaves. Pro_CAT2_:GFP-CAT2 and Pro_CAT2_:CAT2-GFP were co-expressed with the peroxisomal marker Pro_35S_:mCherry-PTS1(SKL). Scale bar: 20 μm. Scanning of green GFP and red mCherry fluorescence intensity through individual cells encompassing two peroxisomes with scanning position and direction indicated by 20 µm red arrows (right panels).

## Notes

### Competing Interest Statement

The authors have declared no competing interest.

